# Identification of Candidate Protein Biomarkers Associated with Domoic Acid Toxicosis in Cerebrospinal Fluid of California Sea Lions (*Zalophus californianus*)

**DOI:** 10.1101/2024.05.03.592242

**Authors:** Gautam Ghosh, Benjamin A. Neely, Alison M. Bland, Emily R. Whitmer, Cara L. Field, Pádraig J. Duignan, Michael G. Janech

## Abstract

Since 1998, California sea lion (*Zalophus californianus*) stranding events associated with domoic acid toxicosis have consistently increased. Outside of direct measurement of DA in bodily fluids at the time of stranding, currently there are no practical non-lethal clinical tests for the diagnosis of domoic acid toxicosis (DAT) that can be utilized in a large-scale rehabilitation facility. Proteomic analysis was conducted to discover candidate protein markers of DAT using cerebrospinal fluid from stranded California sea lions with acute DAT (n = 8), chronic DAT (n = 19), or without DAT (n = 13). A total of 2005 protein families were identified experiment-wide (FDR < 0.01). Of these proteins, 83 were significantly different in abundance across the three groups (adj. p < 0.05). Cytoplasmic malate dehydrogenase (MDH1), 5’-3’ exonuclease PLD3, disintegrin and metalloproteinase domain-containing protein 22 (ADAM22), 14-3-3 protein gamma (YWHAG), neurosecretory protein VGF, and calsyntenin-1 (CLSTN1) were able to discriminate California sea lions with or without DAT (ROC > 0.75). Immunoglobulin kappa light chain-like (IGKV2D-28), receptor-type tyrosine-phosphatase F (PTRPF), kininogen-1 (KNG1), prothrombin (F2), and beta-synuclein (SNCB) were able to discriminate acute DAT from chronic DAT (ROC > 0.75). Interestingly, proteins involved in alpha synuclein deposition were over- represented as classifiers of DAT and many of these proteins have been implicated in a variety of neurodegenerative diseases. These proteins should be considered potential markers for DAT in California sea lions, as well as markers to discriminate between acute or chronic DAT, and should be considered priority for future validation studies as biomarkers. All MS data have been deposited in the ProteomeXchange with identifier PXD041356 (http://proteomecentral.proteomexchange.org/dataset/PXD041356).

## Introduction

In 1998, the first strandings of California sea lions (*Zalophus Californianus;* CSL) associated with domoic acid toxicosis (DAT) were reported.[1] Since then, an average of 108 CSL strandings per year have been reported at a single rehabilitation center in the last two decades (Cara Field, The Marine Mammal Center, 2022, pers. comms.). DAT is caused by the ingestion of domoic acid, a neurotoxin produced by diatoms belonging to the *Pseudo-nitzschiza* genera. It is transferred through trophic levels, with northern anchovies (*Engraulis mordax*) acting as a common vector prey for CSLs.[2][3] In the brain, domoic acid primarily affects the hippocampus, and activates three sub-types of ionotropic receptors: AMPA, NMDA, and kainate receptors[4], which leads to increased intracellular concentration of cytosolic free calcium ions (Ca^2+^), neuronal excitation, manifesting clinically as seizures, leading to neuronal necrosis and, ultimately, hippocampal _atrophy._[5][6][7]

In the absence of empirical data, it is assumed that acute DAT occurs when CSLs consume a high dose of contaminated prey over a short period of time, often in a single event, whereas chronic exposure to domoic acid occurs when CSLs are exposed to the toxin over a prolonged period. Antemortem diagnosis of DAT is challenging. Immediately following exposure, domoic acid can be detected in urine, feces, and milk.[8] However, domoic acid is cleared from the body within days [9][10][11]. In pregnant females, domoic acid may also be detected in amniotic and allantoic fetal fluids.[8] Some clinical pathologic changes such as eosinophilia have been associated with acute domoic DAT.[12] Clinical signs of neurologic disease including obtundation, abnormal movements, impaired vision, and seizures are suggestive, particularly if these are observed in clusters of animals during a known domoic acid-producing algal bloom.[12] However, clinical pathologic changes and clinical signs are nonspecific for DAT, can persist well beyond clearance of domoic acid from the body, and are poor predictors of response to therapy or recovery.[13] Acute or chronic exposure to domoic acid may result in chronic neurologic abnormalities such as epilepsy, leading to impaired foraging and subsequent starvation. Structural neurologic changes including hippocampal atrophy associated with chronic DAT can be detected via magnetic resonance imaging (MRI) or post-mortem brain histology (Figure 1).[14] Adding to the difficulty in diagnosing antemortem DAT in free-living CSLs, it is not possible to know the amount of domoic acid ingested or the course of intoxication.[15]

**Figure 1:**
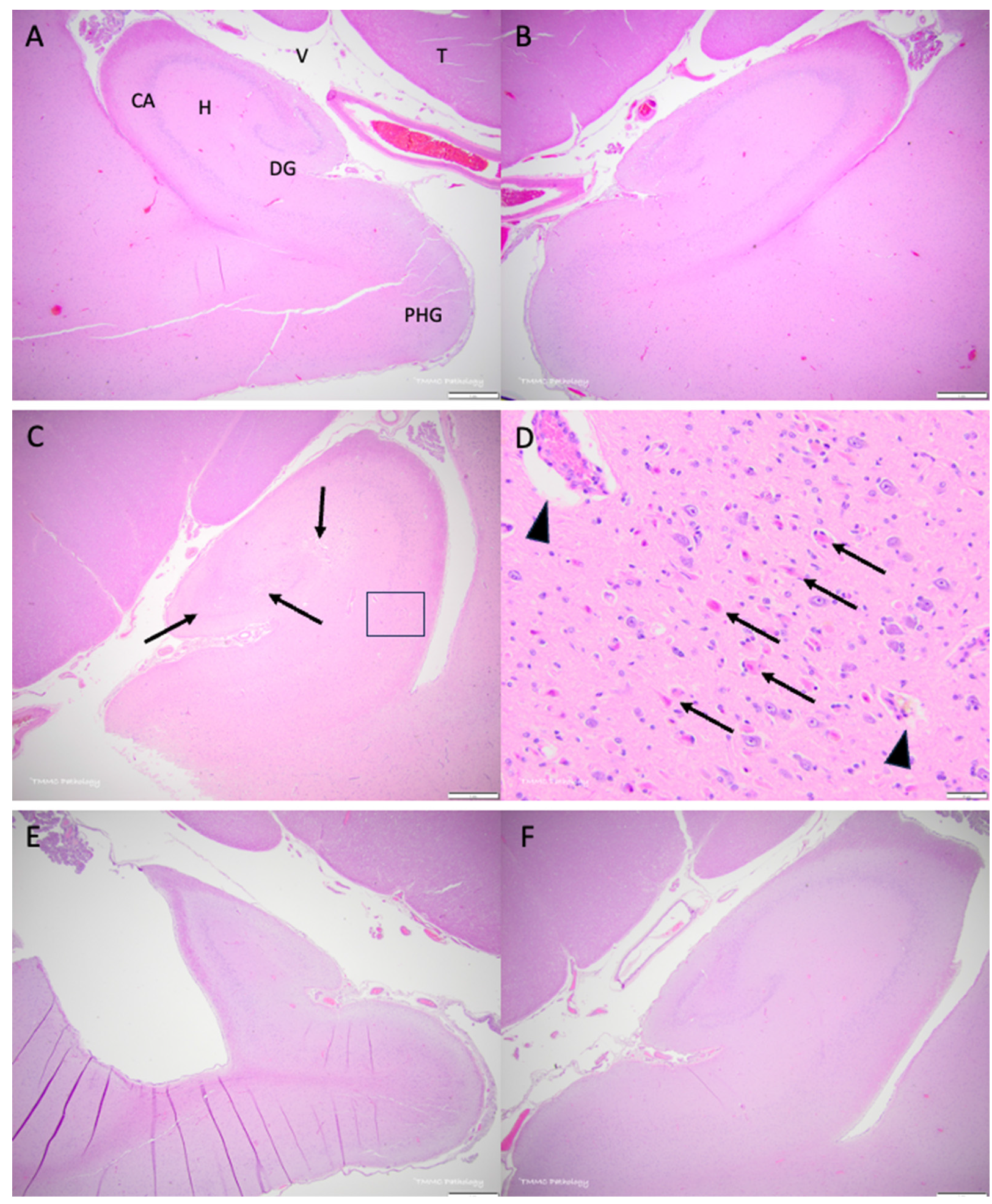
Domoic acid-associated hippocampal histopathology in California sea lions, *Zalophus californianus*. All slides are stained with hematoxylin and eosin and the scale bar is as shown in each panel. (A) Left ventral hippocampal complex from a representative sample without domoic acid pathology. Anatomical features as indicated: ventral hippocampus (H), neurons of the cornu ammonis (CA), neurons of the dentate gyrus (DG), the temporal horn or the lateral ventricle (V), thalamus (T) and parahippocampal gyrus (PHG). (B) Right hippocampal complex from the same individual as A. (C) Representative sample with acute domoic acid toxicity showing hippocampal swelling and edema (pallor in this low power view as indicated by arrow). (D) Higher power view of the area indicated by a box in C showing numerous necrotic neurons (arrows) in the cornu ammonis and perivascular edema (arrowheads). (E) Left hippocampus showing severe atrophy (contraction) of the left hippocampal complex (chronic DAT) with negligible change on the right side of this representative case (F).

Because DAT impacts the brain more than any other part of the body, cerebrospinal fluid (CSF) offers potential as a non-lethal source of biomarkers that could aid in diagnosis and study of DAT.[16][17][18] In the only previous study of CSF in CSLs, Neely et al. examined the protein composition in CSLs with acute DAT, chronic DAT, and absence of DAT (non-DAT).[19] However, this pilot study was limited by the small sample size, an unbalanced design, and the lack of an annotated California sea lion genome from which proteins could be identified. These limitations are likely to have impacted the generalizability of the study results to a larger population. However, despite these shortcomings, results indicated that biomarkers of DAT in CSF should be examined in greater detail using a larger and more rigorously qualified cohort. Recently published and annotated by NCBI RefSeq in 2021, the CSL genome is now available, which allows for a more comprehensive and accurate identification of CSL proteins.[20] The objectives of this study were: 1) to identify candidate protein biomarkers in cerebrospinal fluid from CSLs that can discriminate individuals with DAT from those without DAT, and 2) to identify candidate proteins that can discriminate CSLs with acute DAT from those with chronic DAT.

## Methods

### Sample Collection and Inclusion Criteria

Cerebrospinal fluid samples were collected post-mortem from stranded CSL in a rehabilitation facility (The Marine Mammal Center, TMMC, Sausalito, CA, USA) between 2016 and 2021 under NOAA permit number 18786-04. Individuals were euthanized at the direction of the attending veterinarian due to severe illness or injury with grave prognosis; clinical decision making was independent of study enrollment. Individuals were positioned in lateral recumbency with 90 degrees of neck ventroflexion and CSF was collected aseptically from the spinal canal accessed via the atlanto-occipital joint with a 3.5-inch 18G needle (Quinke spinal needle, Beckton Dickinson Franklin Lakes NJ USA) within 1 hour following euthanasia. The initial 0.5 to 1mL of sample was discarded to reduce contamination, and 2 to 4mL were collected directly into cryovials without additives. Samples were frozen at - 80°C within 12 hours of collection.

DAT status was established by antemortem clinical signs, detection of DA in feces, urine, or milk, gross necropsy and histopathology. Non-DAT individuals had no observed antemortem neurologic abnormalities, primary cause of death as determined by gross necropsy to be inconsistent with DAT (e.g., urogenital carcinoma, pyelonephritis, trauma), and/or no abnormalities detected on brain histology. DAT individuals demonstrated abnormal neurologic status antemortem, DA detected in body fluids, and/or structural abnormalities detected on histopathology. Cases were further classified as having acute or chronic DAT based on histopathology. Additionally, blood samples were collected antemortem, typically within 3 days of admission to the rehabilitation facility, for complete blood count (ABC Plus analyzer, SCIL Vet America, Gurnee IL USA), manual white blood cell differential, and blood chemistry (Axcel clinical chemistry analyzer, Alfa Wasserman-West, Caldwell NJ USA).

The subjects were categorized into non-DAT and DAT based on clinical signs, presence of DA in body fluids, or histologic features of the hippocampal complex. The DAT samples were then further classified as acute DAT and chronic DAT based on disease progression through pathology, and whether the sea lion re-stranded and had a previous DA diagnosis. Included samples (n=40) were collected from adult (25 females, 2 males), sub-adult (3 females, 7 males) and juvenile (3 males) CSLs (Table 1). CSF protein concentration was estimated using the pyrogallol method (QuanTtest, Quantimetrix®, Redondo Beach, CA) and verified using polyacrylamide gel electrophoresis against albumin standards.

**Table 1:** Demographics, hematology and serum biochemistry data for California sea lions from the study. (A denotes P<0.05 versus acute, B denotes P<0.05 versus chronic, Dunn’s post-hoc test). Numbers inside parentheses indicate N.

|  | Acute DAT | Chronic DAT | Non-DAT | P-value<br>(Kruskal Wallis) |
| --- | --- | --- | --- | --- |
| Sex (Male/Female) | (2/6) | (5/14) | (4/9) | 0.37 |
| Age Class |  |  |  |  |
| Juvenile | 0 | 1 | 2 |  |
| Subadult | 1 | 5 | 4 |  |
| Adult | 7 | 13 | 7 |  |
| Complete Blood Count |  |  |  |  |
| WBC (103/μL) | 10.1 ± 0.5 (4) | 11.5 ± 5.6 (13) | 15.9 ± 10.4 (8) | 0.56 |
| Lymphocytes (103/μL) | 16 ± 10 (4) | 23 ± 10 (13) | 19 ± 8 (8) | 0.49 |
| Eosinophils (103/μL) | 8 ± 4 (2) | 5 ± 4 (12) | 3 ± 2 (8) | 0.27 |
| RBC (106/μL) | 4.46 ± 0.34 (4) | 3.95 ± 0.98 (13) | 3.78 ± 0.89 (8) | 0.66 |
| Hematocrit (%) | 45.8 ± 4.2 (4) | 40.9 ± 10.2 (13) | 39.5 ± 11.2 (8) | 0.68 |
| Hemoglobin (g/dL) | 16.8 ± 1.8 (4) | 14.5 ± 3.7 (13) | 13.9 ± 3.7 (8) | 0.51 |
| RDW (%) | 15.1 ± 0.7 (4) | 15.2 ± 0.9 (13) | 16 ± 1.3 (8) | 0.44 |
| MCHC (g/dL) | 36.7 ± 1.4 (4) | 35.5 ± 1.2 (13) | 35.5 ± 1.7 (8) | 0.6 |
| MCH (pg) | 37.7 ± 2.4 (4) | 36.7 ± 2.1 (13) | 36.7 ± 2.8 (8) | 0.84 |
| MCV (fL) | 102 ± 3 (4) | 104 ± 5 (13) | 104 ± 7 (8) | 0.89 |
| Platelets (109/L) | 410 ± 90 (4) | 389 ± 128 (13) | 383 ± 134 (8) | 0.98 |
| MPV (fL) | 8.9 ± 0.6 (4) | 8.9 ± 0.8 (13) | 9 ± 0.9 (8) | 0.93 |
| Serum Chemistry |  |  |  |  |
| Blood urea nitrogen (mg/dL) | 52 ± 77 (5) | 33 ± 29 (15) | 134 ± 101 (9)<sup>A,B</sup> | **0.005** |
| Creatinine (mg/dL) | 1.22 ± 0.88 (5) | 0.87 ± 0.24 (15) | 2.93 ± 2.06 (9) | 0.08 |
| BUN-creatinine ratio | 27.51 ± 21.26 (5) | 38.65 ± 25.7 (15) | 48.61 ± 16.61 (9) | 0.11 |
| Alkaline phosphatase (U/L) | 27 ± 4 (4) | 38 ± 23 (9) | 149 ± 310 (6) | 0.74 |
| Aspartate aminotransferase (U/L) | 23 ± 11 (4) | 28 ± 32 (13) | 112 ± 220 (8) | 0.76 |
| Gamma-glutamyl transferase (U/L) | 120 ± 32 (5) | 227 ± 331 (15) | 205 ± 215 (9) | 0.92 |
| Total bilirubin (mg/dL) | 0.4 ± 0.1 (4) | 0.4 ± 0.2 (13) | 0.5 ± 0.3 (9) | 0.43 |
| Total iron (μg/dL) | 60 ± 24 (4) | 81 ± 41 (12) | 94 ± 49 (6) | 0.53 |
| Glucose (g/dL) | 132 ± 27 (4) | 118 ± 48 (13) | 111 ± 28 (9) | 0.64 |
| Total protein (g/dL) | 7.5 ± 1.1 (4) | 7.8 ± 1.2 (13) | 8.6 ± 1.4 (9) | 0.33 |
| Albumin (g/dL) | 2.7 ± 0.3 (4) | 2.7 ± 0.6 (13) | 2.6 ± 0.7 (9) | 0.95 |
| Globulin (g/dL) | 4.8 ± 0.9 (4) | 5 ± 0.9 (12) | 6 ± 0.9 (9) | 0.1 |
| Sodium (mmol/L) | 148.2 ± 7.4 (5) | 151.2 ± 5.3 (15) | 162.8 ± 15.2<br>(9)<sup>A,B</sup> | **0.02** |
| Phosphorus (mg/dL) | 7.9 ± 3.5 (5) | 6.6 ± 2 (15) | 9.2 ± 3.2 (9) | 0.16 |
| Calcium (mg/dL) | 8.8 ± 0.4 (5) | 8.7 ± 0.8 (15) | 8.9 ± 0.8 (9) | 0.8 |
| Potassium (mmol/L) | 4.99 ± 1.24 (5) | 4.92 ± 1.72 (15) | 4.44 ± 0.47 (9) | 0.89 |
| Creatine kinase (U/L) | 189 ± 85 (4) | 421 ± 418 (13) | 2618 ± 5758 (9) | 0.35 |

Abbreviations: MCH – mean corpuscular hemoglobin, MCHC – mean corpuscular hemoglobin concentration, MCV – mean red cell volume, MPV – mean platelet volume, RBC – total red blood cells, RDW – red cell distribution width, WBC – total white blood cells, BUN – blood urea nitrogen.

### Protein Digestion

The CSF samples were digested in random batches to minimize investigator bias. CSF (100 µg) was mixed with an equal volume of 2 x Lysis Buffer (10% SDS (sodium dodecyl sulfate) (volume fraction), 100 mmol/L TEAB (triethylammonium bicarbonate), pH 7.55), and vortexed. Samples were reduced in a final concentration of 10 mmol/L dithiothreitol (DTT), heated at 60 ℃ for 30 min, and alkylated with a final concentration of 20 mmol/L chloroacetamide (CAA) for 30 min in the dark prior to digesting with 5 µg of Pierce^TM^ trypsin protease (1:20), using S-Trap digestion columns (Protofi). Solid phase extraction was conducted using C18 spin columns (Affinisep^TM^). The samples were then eluted using 0.5 % formic acid (volume fraction) in 50 % acetonitrile (volume fraction), and dried by SpeedVac for three hours, after which the samples were resuspended in 0.1 % formic acid (volume fraction). Tryptic peptides were quantified by a quantitative colorimetric peptide assay (ThermoFisher Scientific^TM^) prior to analysis by mass spectrometry.

### LC-MS/MS

The peptide samples were randomized prior to injection to minimize run bias. The peptide mixtures were separated and analyzed using an UltiMate 3000 Nano LC coupled to a Fusion Lumos Orbitrap mass spectrometer (Thermo Fisher Scientific). One µg of peptide was loaded onto a PepMap 100 C18 trap column (75 µm id x 2 cm length; Thermo Fisher Scientific) at 3 µL/min for 10 min with 2 % acetonitrile (volume fraction) and 0.05 % trifluoroacetic acid (volume fraction) followed by separation on an Acclaim PepMap RSLC 2 µm C18 column (75µm id x 25 cm length; Thermo Fisher Scientific) at 40 °C. Peptides were separated along a 65 min two-step gradient of 5% to 30

% mobile phase B [80 % acetonitrile (volume fraction), 0.08 % formic acid (volume fraction)] over 50 min followed by a ramp to 45 % mobile phase B over 10 min and lastly to 95 % mobile phase B over 5 min, and held at 95 % mobile phase B for 5 min, all at a flow rate of 300 nL/min.

The Thermo Fusion Lumos was operated in positive polarity mode with 30% RF lens, data-dependent mode (topN, 3 s cycle time) with a dynamic exclusion of 60 s (with 10 ppm error). Full scan was set at 60,000 for a mass range of m/z 375 to 1500. Full scan ion target value was approximately 4.0x10^5^ allowing a maximum injection time of 50 ms. An intensity threshold of 2.5x10^4^ was used for precursor selection, including charge states 2 to 6. Data-dependent fragmentation was performed using higher-energy collisional dissociation (HCD) at a normalized collision energy of 32 with quadrupole isolation at m/z 1.3 width. The fragment scan resolution using the orbitrap was set at 15000. The ion target value was 2.0x10^5^ with a 30 ms maximum injection time.

### Data Processing Protocol

Thermo .raw files were searched using Maxquant (v2.0.3.1). The databases that were specified for the search were NCBI Zalophus californianus Annotation Release 101; GCF_009762305.2 (21,397 sequences) and the common Repository of Adventitious Proteins database (cRAP; the Global Proteome Machine) (116 sequences). The search parameters were as follows: trypsin was specified as the enzyme allowing for two mis-cleavages; carbamidomethyl (C) was selected as a fixed modification. Deamidated (NQ), pyro-Glu (n-term Q), and oxidation (M) were selected as variable modifications. The data was visualized using Scaffold (v5.1.1, Proteome Software, Portland, OR, USA) and false discovery rate set to 1 % for peptide and protein identifications. The spectral count data were normalized by arithmetic mean weighting and exported to RStudio (v1.4).

### Data Analysis

CSF protein concentration between California sea lions across the three groups were compared using a one-way ANOVA test. Serum chemistry and hematology data were compared using the Kruskal-Wallis test and, when suitable, the Dunn’s post-hoc test. Area under receiver operator curves (AuROCs) were calculated using pROC script on R (version 4.2.2), to determine classification performance for individual proteins and blood chemistry/hematology data. Medical records were reviewed for medications administered antemortem to each animal. Medications were categorized as non-steroidal anti-inflammatory drugs (NSAID; e.g., carprofen, meloxicam), anti-epileptic drugs (AEDs; e.g., phenobarbital, lorazepam), antibiotics (e.g., cephalexin, ciprofloxacin), gastroprotectants (e.g., famotidine, omeprazole), corticosteroids (e.g., prednisone, dexamethasone), anthelmintics (e.g.,ivermectin, ponazuril), and anti-oxidants (alpha lipoic acid – SQ). The pharmaceutical data were analyzed between groups by Chi-squared analysis. Exponentially modified protein abundance index (emPAI) values for CSF proteins were used to calculate average molar ratios. The average molar ratios of these proteins were used to rank the protein abundances per group.[20] Prior to the statistical analysis, the identified proteins accession numbers were “humanized” using PAW-BLAST (https://github.com/pwilmart), a script that compares protein sequences from one FASTA protein database against another utilizing BLAST tools, to replace CSL NCBI accession numbers with human NCBI accession numbers. Using the humanized accession numbers, g:Profiler (https://biit.cs.ut.ee/gprofiler/gost) was then utilized to investigate the proteins found in the CSF.[21] In g:Profiler, the statistical domain scope was set to “only annotated genes”, significance threshold was set to “g:SCS threshold”, the user threshold was set to “0.05”, and the Numeric IDs were treated as “ENTREZGENE_ACC”. “Human Protein Atlas” was selected under protein database. For the differential analysis of the proteomic data, statistical differences between the proteins across the three groups were calculated using Kruskal-Wallis test followed by Dunn’s post-hoc test when applicable. P-values were adjusted for false-discovery using the Benjamini-Hochberg method.[22] Proteins were considered significant when the adjusted p-value was less than 0.05. Individual candidate markers were assessed for statistical performance, and AuROCs were further utilized to rank candidate markers.

## Results

### Study Population

CSF samples (n = 40) were analyzed from adult (25 females, 2 males), sub-adult (3 females, 7 males) and juvenile (3 males) CSL (Table 1). Serum chemistry and hematology data were available for 5 acute DAT individuals, 15 chronic DAT individuals and 9 non-DAT individuals. Within this subset, cases diagnosed with both acute and chronic DAT had significantly lower levels of blood urea nitrogen (BUN) and sodium compared to the individuals with DAT (Table 1, p < 0.05, Kruskal-Wallis test, Dunn’s post-hoc test). BUN was 1.6-fold lower in individuals with acute DAT, and 3-fold lower in individuals with chronic DAT. Serum sodium levels were 0.1-fold lower in individuals with acute DAT, and 0.08-fold lower in individuals with chronic DAT. AuROC analysis showed that most hematology parameters lacked sensitivity and specificity for the diagnosis of DAT in CSLs (Supplemental Table 1), although this only reflects a subset of individuals that had clinical data and conclusions were not drawn from this information. The analysis of the pharmaceutical regimen for the CSLs (Supplemental Table 2) showed that 38 % more individuals without DAT were treated with anti-inflammatory non-NSAIDs than individuals with acute DAT (p < 0.05, Chi-squared test), and 44 % more individuals with acute DAT were treated with anti-seizure drugs than individuals without DAT (p < 0.05, Chi-squared test).

### CSF Proteome

There was no difference (One-way ANOVA test, p = 0.368) in mean CSF total protein concentration between sea lions in the acute DAT group (0.93 ± 0.37 μg/μL), chronic DAT group (0.68 ± 0.24 μg/μL), or the non-DAT group (0.78 ± 0.29 μg/μL). A total of 2005 proteins were identified experiment-wide (FDR < 0.1). Protein ranking by average molar ratio of the 20 most abundant proteins showed consistency amongst the highest abundance proteins across all groups (Figure 2). Albumin and Transthyretin were the two most abundant proteins in California sea lion CSF for all groups, together totaling approximately 50 % of protein composition ratio. Sixteen of the top 20 abundant proteins were common across the three groups, which showed that the protein composition of the CSF was not drastically altered by neurotoxicity. Search of the Human Protein Atlas database through g:Profiler[23] indicated that 79.04 % of proteins in the Non-DAT CSLs, 77.85 % of proteins in the Acute DAT CSLs, and 91.23 % of proteins in the Chronic DAT CSLs were associated with the cerebral cortex (Adjusted p-values: Non-DAT = 0.012, Acute DAT = 4.18x10^-58^, Chronic DAT = 0.00016).

**Figure 2:**
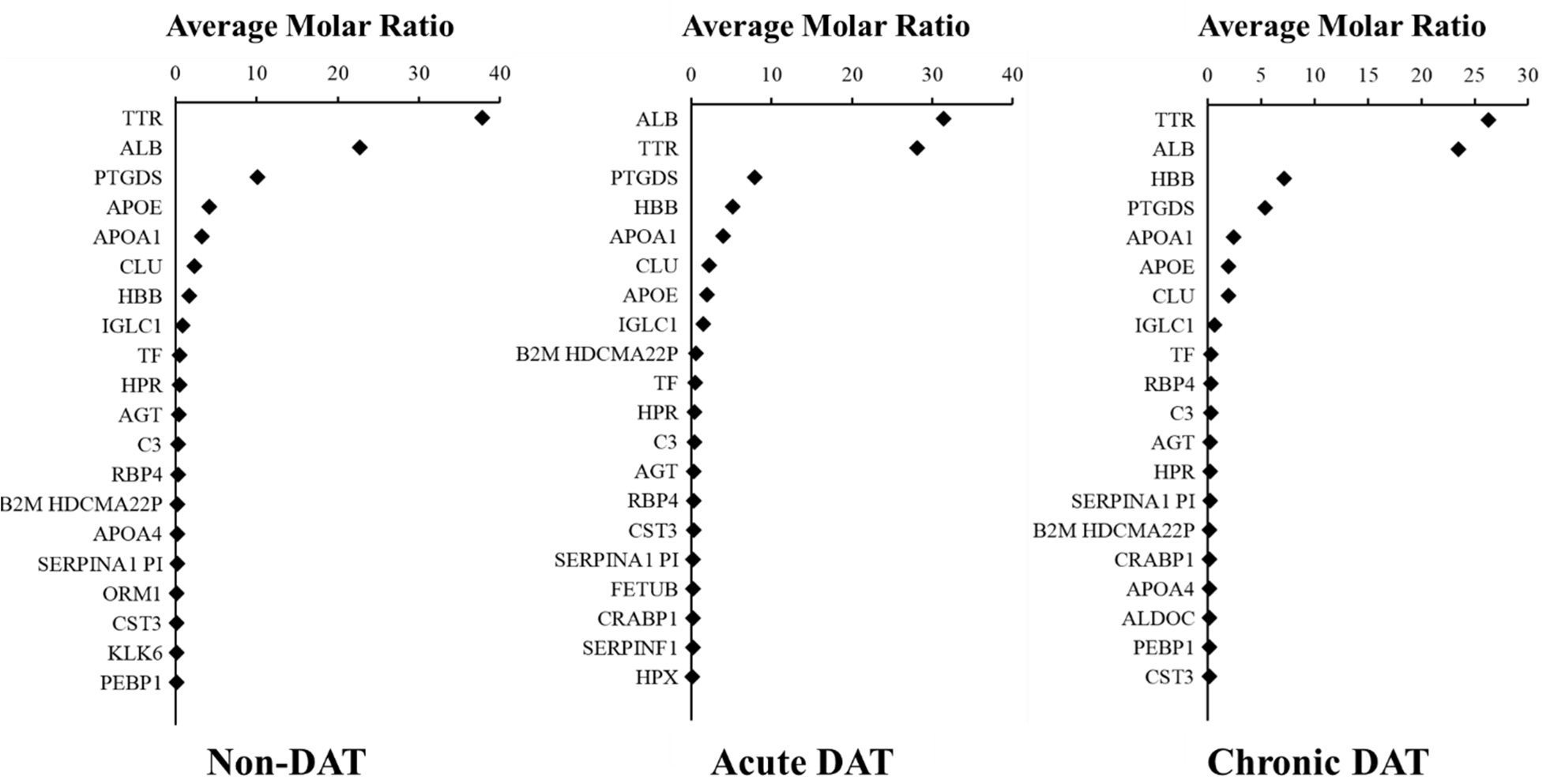
Ranked average molar ratio (%) for CSF proteins, determined using normalized emPAI values, for Acute DAT, Chronic DAT and Non-DAT. The top 20 of the most abundant proteins are shown. Gene symbols are assigned to each of their respective proteins. 16 of the top 20 abundant proteins were common across the three groups.

### Differential Analysis

Among all three groups of CSLs, 83 proteins were significantly different (p < 0.05, Kruskal-Wallis, Benjamini-Hochberg corrected, Supplemental Table 3). Post-hoc analysis of significant proteins revealed 24 of the 83 proteins were significantly different between acute DAT and chronic DAT, 31 proteins were significantly different between acute DAT and non-DAT, and 32 proteins were significantly different between chronic DAT and non-DAT (p<0.05, Dunn’s post-hoc, Benjamini-Hochberg corrected). The top 10 proteins with the lowest p-values across the three groups were beta-synuclein (SNCB), immunoglobulin kappa light chain-like (IGKV2D-28), receptor-type tyrosine-protein phosphatase F (PTRPF), 5’-3’ exonuclease PLD3 (PLD3), cytoplasmic malate dehydrogenase (MDH1), microtubule-associated protein 2 (MAP2), 14-3-3 protein gamma (YWHAG), neurosecretory protein VGF (VGF), disintegrin and metalloproteinase domain-containing protein 22 (ADAM22), brain acid soluble protein 1 (BASP1).

### Area under Receiver Operator Curves (AuROC)

Sixty-eight proteins had an AuROC greater than 0.7 between CSLs with DAT and CSL without DAT (Figure 3). Of these 68 proteins, three proteins had an AuROC greater than 0.80: 5’-3’ exonuclease PLD3 (PLD3, AuROC=0.84), Disintegrin and metalloproteinase domain-containing protein 22 (ADAM22, AuROC=0.82), and 14-3-3 protein gamma (YWHAG, AuROC=0.801) (Supplemental Table 1). Similarly, sixty-six proteins had an AuROC greater than 0.7 between samples with acute DAT and samples with chronic DAT (Figure 3), of which 11 proteins had an AuROC greater than 0.8, and one protein had an AuROC greater than 0.9: immunoglobulin kappa light chain-like (IGKV2D-28, AuROC=0.901) (Supplemental Table 4).

**Figure 3:**
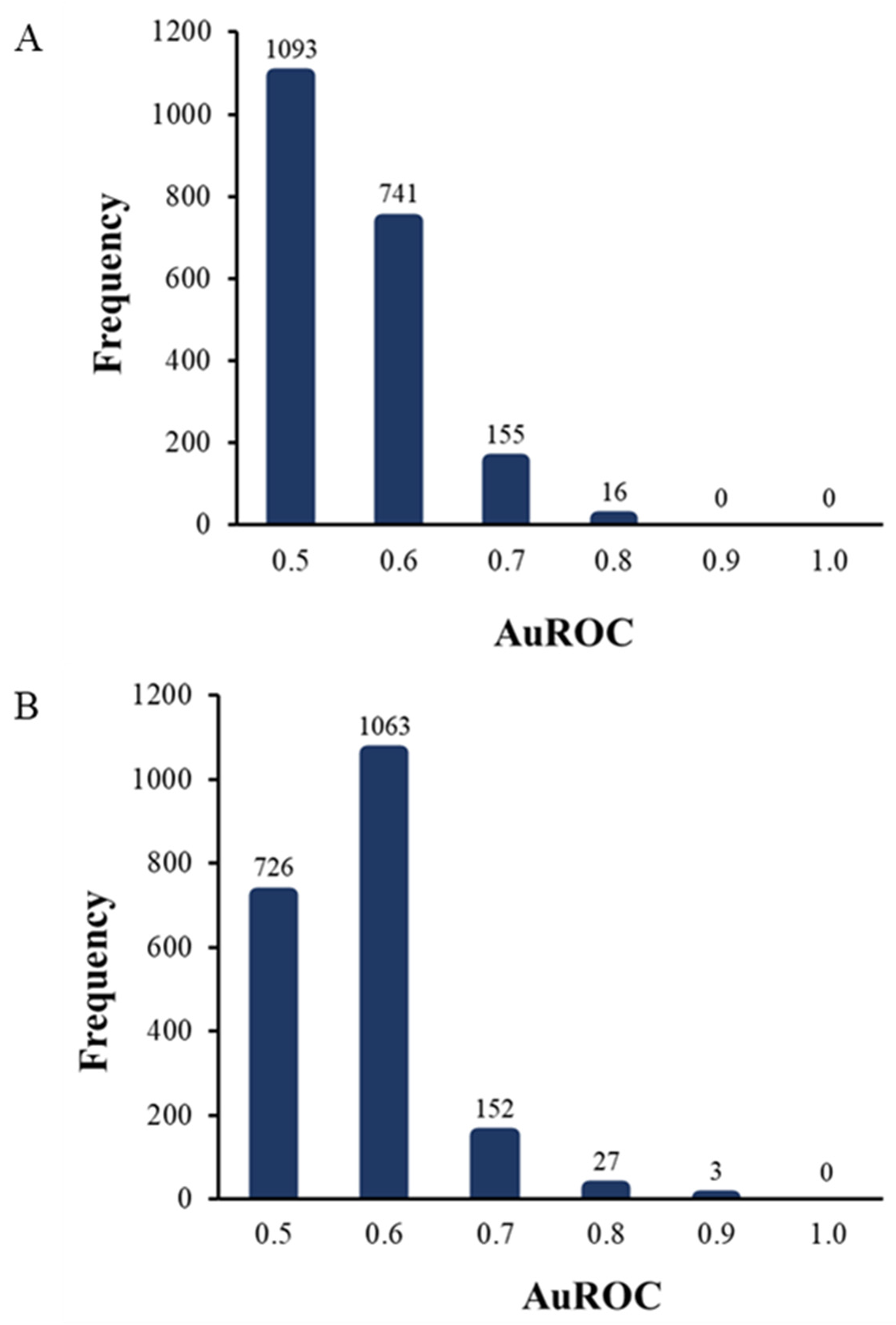
(A) Area under the ROC curve (AuROC) frequency distribution of individual proteins for non-DAT versus all DAT. Receiver operator characteristic curves were constructed for each protein using weighted spectral count data. AuROC values were binned at every 0.1 ± 0.05. **(B)** Area under the ROC curve (AuROC) frequency distribution of individual proteins for acute DAT versus chronic DAT. Receiver operator characteristic curves were constructed for each protein using weighted spectral count data. AuROC values were binned at every 0.1 ± 0.05.

The top performing classifier proteins (p<0.05, Kruskal-Wallis Test, Benjamini-Hochberg corrected) with the highest AuROCs in the non-DAT vs DAT comparison were: PLD3 (Acute, - 2.0 fold; Chronic, -2.7 fold versus non-DAT), YWHAG (Acute, 5 fold; Chronic, 2.8 fold versus non-DAT), ADAM22 (Acute, -2.6 fold; Chronic, -2.4 fold versus non-DAT), CLSTN1 (Acute, - 2.2 fold; Chronic, -2.8-fold versus non-DAT), APLP2 (Acute, not different; Chronic, -1.7 fold versus non-DAT), and VGF (Acute, not different; Chronic, -2.5 fold versus non-DAT) (Figure 4, Figure 6, Supplemental Table 4).

**Figure 4:**
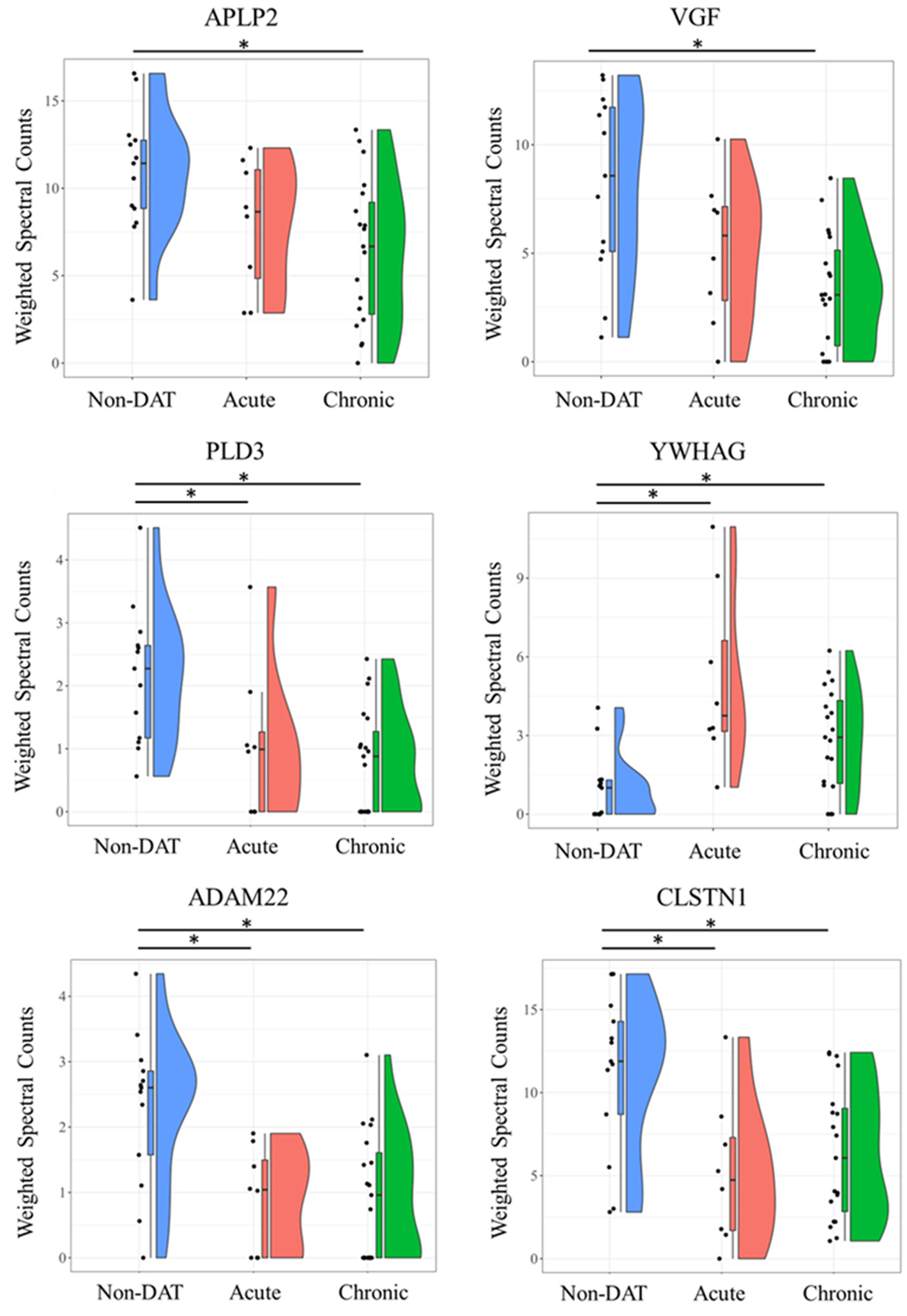
Violin plots showing comparisons of weighted spectra for all individuals plotted across all groups for proteins with highest AUCs between CSLs diagnosed with DAT and Non-DAT: APLP2, VGF, PLD3, YWHAG, ADAM22, and CLSTN1. Proteins across groups were considered different by Kruskal-Wallis test (p<0.05). *denotes significant difference in protein abundance between specific groups (p<0.05, Kruskal-Wallis test, Dunn’s post-hoc test).

The top performing classifiers proteins (p<0.05, Kruskal-Wallis Test, Benjamini-Hochberg corrected) with the highest AuROCs in the chronic DAT vs acute DAT comparison were: IGKV2D-28 (Acute 2.2 fold versus Chronic), PTPRF (not present in Acute), KNG1 (Acute 1.7 fold versus Chronic), F2 (Acute 2.0 fold versus Chronic), IGKV2-29 (Acute 2.1 fold versus Chronic), LGB2 (Acute 2.5 fold versus Chronic) (Figure 5, Supplemental Table 5).

**Figure 5:**
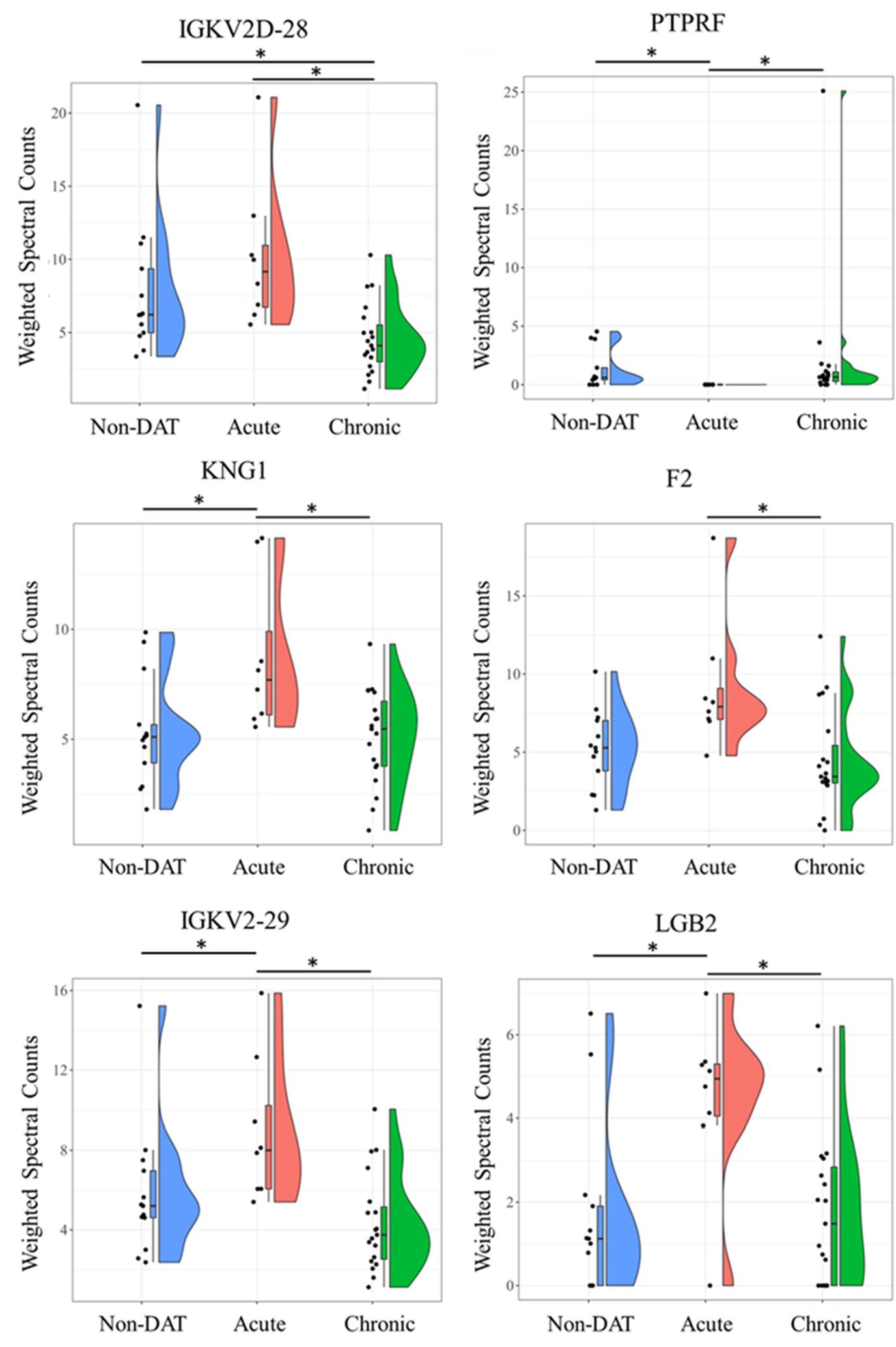
Violin plots showing comparisons of weighted spectra for all individuals plotted across all groups for proteins with highest AUCs between CSLs diagnosed with Acute DAT and Chronic DAT: IGKV2D-28, PTPRF, KNG1, F2, IGKV2-29 and LGB2. Proteins across groups were considered different by Kruskal-Wallis test (p<0.05). *denotes a significant difference in protein abundance between specific groups (p<0.05, Dunn’s post-hoc test).

**Figure 6:**
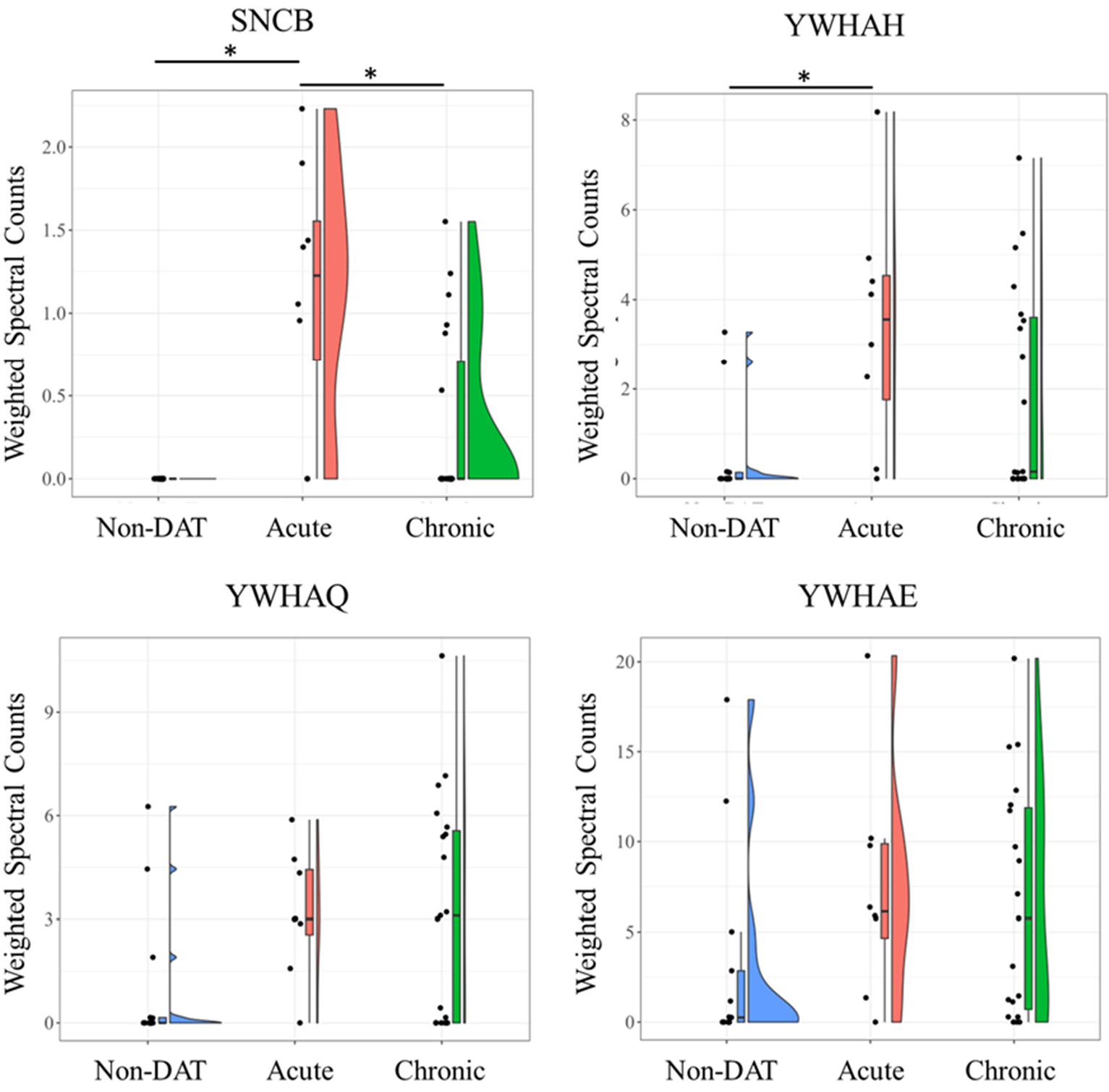
Violin plots showing comparisons of weighted spectra for all individuals plotted across all groups for SNCB, YWHAH, YWHAQ, and YWHAE. Proteins across groups were considered different by Kruskal-Wallis test (p<0.05). *denotes a significant difference in protein abundance between specific groups (p<0.05, Dunn’s post-hoc test).

## Discussion

The purpose of this study was to identify biomarkers in CSF samples for DAT, and to further discover candidate markers in CSF samples to classify California sea lions with acute DAT from chronic DAT. While serum and plasma biomarker studies are more common than CSF biomarker studies, multiple studies have used CSF to discover biomarkers for neurodegenerative disorders in humans.[24][25][26][27][28] Neely et al. (2015) is the only other study of CSF proteins for CSLs with domoic acid intoxication; although, these authors reported several limitations, including the small sample size, disproportionate sex ratios amongst the groups due to availability, and the lack of an annotated CSL genome from which proteins could be identified. Since then, the CSL genome was completed and annotated[29], allowing for more accurate protein identification for peptide spectral matching, and technological advances in mass spectrometry have allowed for a more comprehensive inspection of the proteome. As a result, a total of 2005 proteins were identified in this study, which is 10-fold more than Neely et al. The latter authors only identified 8 proteins as potential classifiers for DAT whereas in our study, 83 proteins were identified to be significantly different experiment wide: 24 significantly different proteins between acute DAT and chronic DAT, 31 significantly different proteins between acute DAT and non-DAT, and 32 significantly different proteins between chronic DAT and non-DAT.

Characteristic of neurodegenerative diseases that result in seizures, albumin and total protein concentration is typically elevated.[30] Although albumin was one of the most abundant proteins in all CSLs, we found no evidence for a significant elevation in CSF albumin or total protein. The lack of difference in albumin or total protein could be explained by timing and therapeutic intervention. The time at which a CSF sample was taken following the last seizure event is unknown and most CSLs with DAT were treated with antiseizure medications as well as some (31%) of the non-DAT CSLs. Timing and intervention were not intentionally balanced for this study therefore conclusions regarding CSF protein abundance require further study.

### Non-DAT vs. DAT Candidate Markers

Sixty-eight CSF proteins were good classifiers of DAT when all acute and chronic animals were grouped together to create a general category. However, some of the classification power of proteins such as VGF nerve growth factor, was largely driven by chronic DAT CSLs because acute DAT CSLs were no different than non-DAT CSLs. This highlights an issue with grouping acute and chronic DAT CSLs together and may partially underline why markers for DAT in CSLs have been difficult to identify.

On the other hand, some proteins that appeared to be good general classifiers of DAT maintain discriminatory power despite the grouping (Figure 4). Of these proteins, phospholipase D family member 3, disintegrin and metalloproteinase domain-containing protein 22, and calsyntenin-1 were lower in CSLs with DAT. The downregulation or loss of function through mutation of these genes have been implicated in neuronal mechanisms of disease progression. ADAM22 is catalytically dormant until it binds to LGI1, a neuronal glycoprotein and this LGI1-ADAM22 complex is important for synapse function and maturation in the post-synaptic membrane.[31][32]**^33]^**Loss of this complex limits AMPA receptor function and results in epileptic signs in a rodent model.[34]

Mutations in the 5’-3’ exonuclease PLD3 gene, also known as phospholipase D3, have previously been attributed to patients with late-onset Alzheimer’s Disease.[35][36][37] Loss of function leads to increased levels of amyloid beta, which accumulates in the brains of patients with the disease.[38]

Calsyntenins bind calcium and play an important role in the production of the amyloid-β peptide by regulating the axonal transport of the amyloid precursor protein.[39] The downregulation of calsyntenin-1 leads to a disruption of axonal transport, which occurs during Alzheimer’s disease. The downregulation of calsyntenin-1 has also been observed in patients with frontotemporal dementia and this protein has been identified as a potential biomarker in CSF for neurodegenerative disorders.[40][41][42]

### Acute DAT vs. Chronic DAT Candidate Markers

The diagnosis of acute versus chronic DAT is difficult to make by clinical observation alone and the definitive characterization requires post-mortem histopathology or antemortem MRI. No CSF proteins were perfect classifiers of acute or chronic DAT; however, there were twelve reasonable candidates with AUC above 0.80 (Table S1, Figure 5). Two immunoglobulin light chains (IGKV2D-28 and IGKV2-29) were of the highest performing classifiers (AUC > 0.88) and have similar expression profiles where CSLs with acute DAT have on average two-fold higher levels than chronic DAT animals. The elevation in light chains in the acute group may reflect intrathecal production of immunoglobulins as described in human patients with epilepsy[43], or impairment in the blood brain barrier[44]; whereas the reduction in light chains in the chronic group relative to the acute group could be due to the course of antiseizure therapy[45], or resolution of inflammatory changes in the more chronic cases as observed on histology, or both.

The most striking difference between acute and chronic DAT animals involves receptor-type tyrosine-protein phosphatase F (PTPRF, Figure 5). Receptor-type tyrosine-protein phosphatase F, also known as Leukocyte common antigen-related phosphatase (LAR), is a tyrosine phosphatase expressed in neurons and microglia of the brain[46]. Receptor-type tyrosine phosphatases, including PTPRF, play a vital role in signal transduction pathways, synaptogenesis, neurogenesis, blood-brain barrier maintenance and cell cycle regulation.[47][48] PTPRF knock out mice display reduced innervation of the hippocampus, an impairment in spatial learning, and hyperactivity.[49] The reduction PTPRF may represent phenotypic progression of the disease or could represent a protective mechanism to reduce NMDA signaling during excitotoxic injury involving overstimulation of NMDA receptors which is one of the receptor targets of domoic acid.[50]

### Synucleinopathy Proteins and DAT

One of the more striking findings in this study involves the differential abundance of beta-synuclein and 14-3-3 proteins. Beta-synuclein was only identified in CSF of CSLs with DAT (Figure 5). Although the chronic DAT CSL group was not significantly different from the non- DAT group, the presence or absence of peptides in some animals within the chronic DAT group suggest beta-synuclein abundace may be an indicator of acute domoic acid toxicosis and represent a continuum from acute to chronic DAT. Beta-synuclein is a member of the “synuclein” family, that also include alpha-synuclein and gamma-synuclein., and is concentrated in pre-synaptic terminals.[51] Beta-synuclein plays a key role in synucleinopathies, a group of neurodegenerative diseases, where alpha-synuclein misfolds, aggregates to form fibrils, and leads to neuroinflammation and neurotoxicity.[52] Typically associated with Lewy body formation in Parkinsons’s disease, elevated levels of secreted alpha synuclein have been reported in cases of epilepsy which is similar to domoic acid-induced temporal lobe epilepsy in CSLs.[9][53] Elevation of alpha synuclein in epilepsy is further supported by a proteomic study using a pilocarpine mouse model where alpha synuclein was higher in the dentate gyrus of mice with induced seizures.[54] Although no significant differences were noted in the abundance of alpha-synuclein across the three groups, the drastic elevation in beta synuclein and precedence of alpha synuclein from previous reports in humans and mice suggests that alpha synuclein may be choreographing the response observed in CSLs with DAT. Beta-synuclein has been found to have neuroprotective properties that lead to the reduction of alpha-synuclein expression, and binds with high affinity to alpha synuclein monomers to prevent aggregation.[51][55] Recently, CSF and serum beta-synuclein has been implicated as an important biomarker for Alzheimer’s disease associated with synaptic degeneration.[56][57][58][59][60] Interestingly, synucleopathies have not been implicated in domoic acid toxicosis, despite similarities with temporal lobe epilepsy, suggesting this should be a new direction for investigation into mechanisms of progressive neurotoxicosis in CSLs.

14-3-3 proteins are a chaperone protein family that are homologous but functionally diverse, and are categorized into seven isoforms.[61] The 14-3-3 proteins are some of the most abundantly expressed proteins that have been found in the central nervous system, predominantly in the cerebral cortex of the brain.[62][63] These proteins are known to regulate signal transduction, neuronal development, neuroprotection, and cellular processes.[64] The roles of 14-3-3 proteins have previously been studied in kainic acid-induced systems, as well as in neurodegenerative disorders.[65][66][67] Elevated levels of 14-3-3 proteins were discovered in kainic acid-induced rat models, and a similar trend was also observed in human patients with neurodegenerative disorders such as Alzheimer’s disease and Creutzfeldt-Jakob disease.[65][68][69][70]. When DAT sea lions are examined as a combined group 14-3-3 proteins gamma, eta, theta and epsilon were elevated more than two-fold compared to the non-DAT group (Figure 4 and Figure 6). The 14-3-3 proteins are linked to synucleinopathy probably because 14-3-3 protein share physical and functional homology with beta-synuclein and are capable of binding to and inhibiting alpha-synuclein aggregation during synucleinopathic progression.[71][72] In addition to being a candidate biomarker for DAT, 14-3-3 proteins further support alpha synuclein as a node in this CSF proteomic network.

## Conclusion

Of the most promising markers, six CSF proteins were identified as the highest classifiers to distinguish between any DAT and non-DAT individuals. The results of this study also provided a list of five candidate protein markers that should be considered as classifiers of acute or chronic domoic acid intoxication in California sea lions. Interestingly, beta-synuclein was identified as a high classifier for both DAT vs. non-DAT and acute DAT vs. chronic DAT comparisons. The identification of proteins related to synucleinopathies is a new finding for DAT and should be considered in future investigations.

## Data Availability

The proteomics data obtained through mass spectrometry have been archived in the ProteomeXchange Consortium (http://proteomecentral.proteomexchange.org). The dataset is identified as PXD041356 and is available through the PRIDE partner repository.

## Abbreviations

DAT: domoic acid toxicosis
CSF: cerebrospinal fluid
CSL: California sea lion
BUN: blood urea nitrogen

## Acknowledgements

Funding for this project was provided in part through the College of Charleston, Department of Biology Research Fund and storage equipment was purchased with a gift through the Spaulding- Paolozzi Foundation. We would like to thank the Grice Marine Lab for providing laboratory space. We gratefully acknowledge the contributions of Barbie Halaska, Jackie Isbel, Margaret Martinez, Carlos Rios, Jesierose Poblacion, and Mariah Tengler. The identification of certain commercial equipment, instruments, software, or materials does not imply the recommendation or endorsement by the National Institute of Standards and Technology, nor does it imply that the products identified are necessarily the best available for their purpose.

